# Bacterial microcompartment utilisation in the human commensal *Escherichia coli* Nissle 1917

**DOI:** 10.1101/2024.05.13.593892

**Authors:** Chania Clare, Jack W. Rutter, Alex J.H. Fedorec, Stefanie Frank, Chris P. Barnes

## Abstract

Bacterial microcompartments (BMCs) are self-assembled protein structures often utilised by bacteria as a modular metabolic unit, enabling the catalysis and utilisation of less common carbon and nitrogen sources within a self-contained compartment. The *ethanolamine (EA) utilisation (eut)* BMC has been widely demonstrated in enteropathogens, such as *Salmonella enterica*, and current research is exploring its activity in the commensal species that populate the human gut. *Escherichia coli* Nissle 1917 (EcN) is a strong coloniser and probiotic in gut microbial communities, and has been used extensively for microbiome engineering. In this study, the utilisation of ethanolamine as a sole carbon source and the formation of the *eut* BMC in EcN were demonstrated through growth assays and visualisation with transmission electron microscopy. Subsequently, flux balance analysis was used to further investigate the metabolic activity of this pathway. It was found that not only is the utilisation of the *eut* BMC for the degradation of EA as a carbon source in EcN comparable to that of *Salmonella enterica*, but also that ammonium is released into solution as a byproduct in EcN but not in *S. enterica*. Control of EA-dependent growth was demonstrated using different concentrations of the operon inducer, vitamin-B_12_. We show that vitamin B_12_-dependent EA utilisation as the sole carbon source enables growth in EcN, and demonstrate the concurrent formation of the BMC shell and inducible control of the *eut* operon.

## 1 Introduction

The human gut comprises a complex interplay of microorganisms, metabolites and enzymes, where dysbiosis (disruption in the healthy state of the microbiota) is linked to a multitude of health problems [1, 2, 3, 4]. The ecology of this community is complex and can be difficult to restore when perturbed. For example, under specific circumstances a commensal strain of bacteria may become an opportunistic pathogen, which requires the administration of antibiotics to remove [5]. As many antibiotics are non-specific, they can have widespread detrimental effects throughout the gut microbial community, as well as contributing to the increasing prevalence of antimicrobial resistance [6]. Therefore, to understand how to control and treat dysbiosis in a more nuanced predictable manner, it is important to understand not only the colonisation mechanisms of these bacteria, but also their interactions within the gut environment.

*Escherichia coli* Nissle 1917 (EcN) plays an important role preventing pathogen driven dysbiosis, where EcN is commonly used as a probiotic and target of bioengineering to promote and maintain a diverse and stable gut microbial population [7, 8, 9]. EcN both directly and indirectly antagonises multiple enteropathogens through competition and in preventing systemic circulation [10, 11], as well as reducing the invasion capacity of Shiga-like toxin *E. coli, Listeria monocytogenes, Salmonella enteritidis* and *Legionella pneumophila* [12, 13, 14]. Additionally, EcN elicits anti-inflammatory and immune responses [15, 16], as well as contributing to the healthy function of the intestinal epithelial barrier [17].

Bacterial microcompartments (BMCs) encapsulate enzymatic pathways in self-assembling protein shells, and can be found across 23 bacterial phyla, expanding their metabolic potential [18, 19, 20]. The compartment can be beneficial in preventing the exposure of the cell to toxic intermediates, such as aldehydes by coupling its sink reactions to the recycling of an exclusive cofactor pool [21, 22, 23, 24], and in increasing the rate of reaction through the high concentration of enzymes [25]. Many examples of BMC-associated metabolic pathways are present in the human gut [26, 27], where they enable the catabolism of alternative substrates. They commonly confer competitive advantage to enteric pathogens when more typical substrates like glucose are restricted, enabling them to overcome colonisation resistance and rapidly populate the microbiota [28].

For example, in *Salmonella enterica* the capacity for EA catabolism enables the bacteria to sense different gut environments, and therefore enact optimal proliferation strategies, and to interact with the gut environment in order to outcompete the other resident bacteria [29]. In this way, not only does *S. enterica* utilise EA to increase the range of nutrients it can survive on and thus gain a competitive advantage, but is also uses this pathway to change the gut environment for its advantage and adapt to it itself. In anaerobic conditions, EA does not support significant growth through fermentation, however *S. enterica* induces the gut to produce tetrathionate as an alternative electron acceptor, enabling it to utilise EA even in anaerobic conditions [30, 31]. It is held that *S. enterica* uses this niche to rapidly proliferate and cause infection and dysbiosis in the human gut [32, 33]. However, this report looks to investigate whether the human gut commensal *E. coli* Nissle 1917 can also utilise this niche, and perhaps also the gut environment. This could reveal a point of competition that can be exploited for methods of bacterial community control.

This study focuses on the *eut* BMC, which contains the enzymes necessary for the ethanolamine utilisation pathway [20] (Figure 1A). Ethanolamine is a product of phosphatidylethanolamine degradation, a common lipid component of plant and animal cell membranes [34]. Due to the high epithelial turnover in the gut due to mechanical and chemical stress, EA is a naturally abundant carbon source [35]. The *eut* BMC takes up ethanolamine, where - via the toxic and volatile intermediate acetaldehyde - it is catabolised into ethanol, acetyl-phosphate and acetyl-CoA, which is then sequestered into the TCA cycle and contributes to cell growth [36]. In this way, some bacteria can utilise ethanolamine as an alternative carbon and/or nitrogen source when nutrient limited [33].

**Figure 1.**
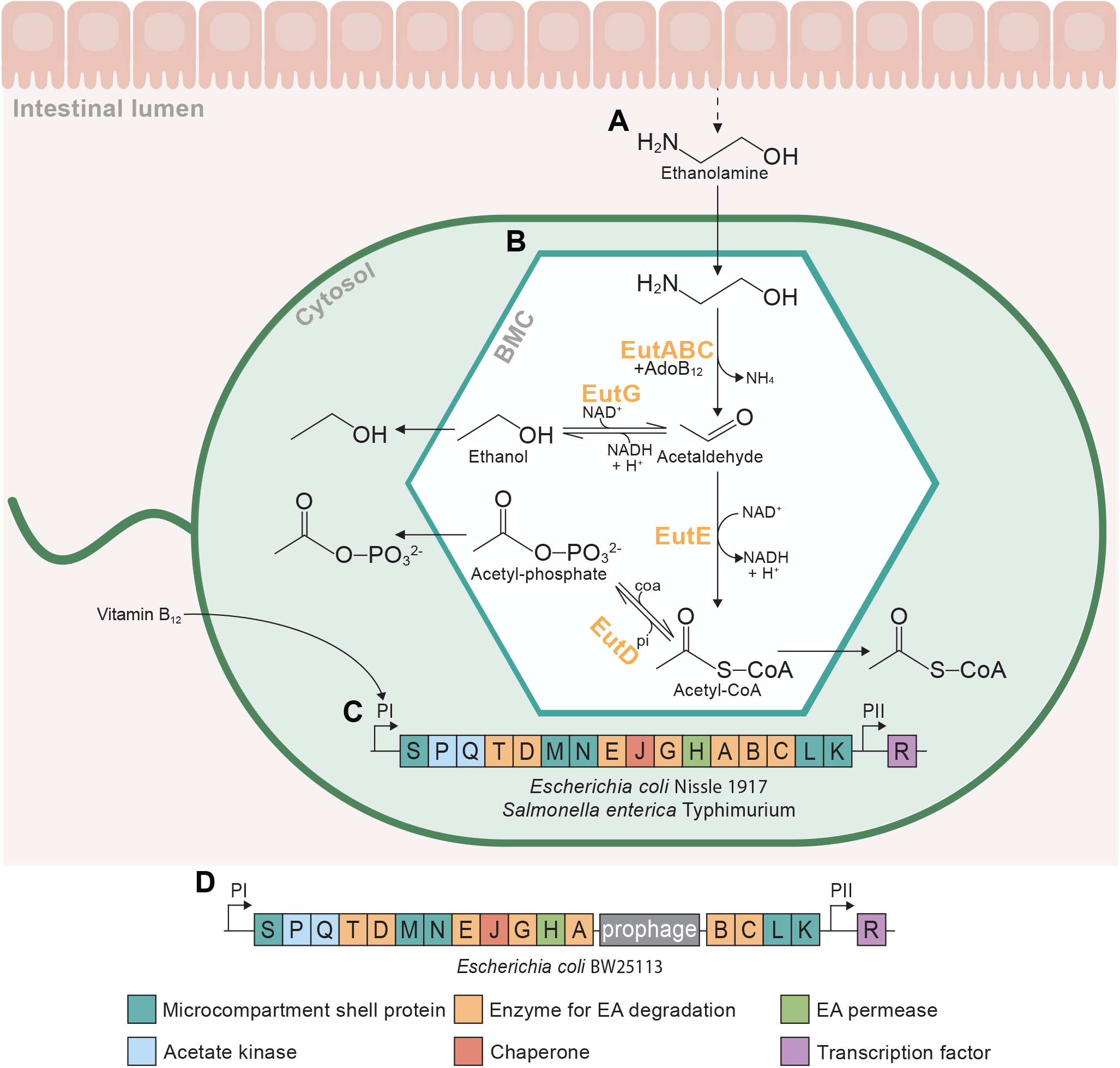
Schematic of the catabolism of ethanolamine by the ethanolamine utilisation (eut) pathway within the bacterial microcompartment (BMC) of *E. coli* Nissle 1917. (**A**) The production of ethanolamine (EA) from lipid membrane degradation in the intestinal lumen. (**B**) EA enters the micro-compartment and is metabolised to ethanol, acetyl-phosphate, and acetyl-CoA. The enclosed environment of the *eut* BMC prevents dissipation of the volatile and toxic intermediate acetaldehyde into the cytosol. Metabolic enzymes encapsulated within the BMC include EA ammonia lyase (EAL, EutBC), EAL reactivase (EutA), alcohol dehydrogenase (EutG), aldehyde dehydrogenase (EutE), and phosphotransacetylase (EutD). (**C**) The *eut* operon in *E. coli* Nissle 1917 and *S. enterica*. Transcription is induced from promoter PI in the presence of both EA and vitamin B_12_, and promoter PII regulates weak constitutive expression of the transcription factor EutR [30]. (**D**) The *eut* operon in *E. coli* BW25113 is interrupted by the CPZ-55 prophage encoding 9 genes translated in the opposing direction between EutA and EutB, preventing the functional expression of the *eut* operon [31, 32].

While the *eut* operon is present in 10.7% of bacteria common to the human gut [27], not all of these have the capacity for the expression and formation of the BMC, due to genetic mutations and prophage interruptions [37]. Many commensal strains of *E. coli* were thought not to have the capacity for EA degradation, perhaps because of the prophage insertion in the common lab strain K12 from which the *E. coli* reference genome is based [38]. A recent study from de Gouveia et al. provides an exciting new overview of the widespread capacity for the degradation of EA as a nitrogen source among 35 *E. coli* human gut commensals, and as a carbon source among three [39]. While EcN has been demonstrated to utilise EA as a nitrogen source [40], it had not yet been demonstrated to utilise EA as a sole carbon or energy source.

The growth rate of a bacterium depends on its ability to successfully occupy its ecological niche, and in the dynamic environment of the gut microbiota, adaptation to varying nutrient availability confers a significant advantage [41, 42]. Many enteropathogens including strains of *E. coli, Salmonella enterica* and *Enterococcus faecalis* have been shown to utilise the *eut* BMC to catabolise EA as an alternative carbon and nitrogen source when the availability of glucose is low to gain a competitive advantage [33, 32, 36, 43]. Much of the previous research into BMC-dependent growth has focused on its role in pathogen colonisation [32, 44, 18, 45, 46], but recent studies have found its presence in gut commensal species too, and call for a broader research focus in the field [47, 39], perhaps demonstrating further complexity in gut bacterial community dynamics than previously thought. In this way, BMCs could be utilised as a tool in microbiome engineering to control the occupation of ecological niches and therefore the growth rates of select bacteria. Further understanding of how BMCs play a role in bacterial population dynamics could enable greater engineering potential of microbial consortia, and control of enteric pathogens.

This study demonstrates the utilisation of EA as a sole energy source in the human gut commensal EcN through the expression and utilisation of the *eut* BMC. The capacity of EcN for EA degradation as a carbon source was shown through growth assays, and the formation of the BMC structure through TEM. The expression of key genes encoding the signature enzyme, shell protein and regulatory protein under conditions of EA utilisation was confirmed with qPCR. Additionally, flux balance analysis was used to make predictions about changes to metabolic output when catabolising EA, which were tested experimentally through ammonia and acetate assays. Finally, the control of *eut* BMC-dependent growth by varying the induction of the native promoter was shown in EcN and *S. enterica*. These results contribute to the understanding of the complex microbial interactions within the human gut.

## 2 Results

### 2.1 EA utilisation in EcN

The capacity of EcN to catabolise EA as the sole carbon source to support cell growth was explored through growth assays under either glucose or EA and vitamin B_12_ supplementation. *Escherichia coli* BW25113 (EcBW), which contains the *eut* operon but with a prophage insertion preventing its functional expression [48] (Figure 1B), was used as a negative control to demonstrate growth under these conditions with the absence of the *eut* BMC and its signature enzymes. *S. enterica*, which has previously been demonstrated to express the genes of the *eut* operon and utilise the associated BMC under these conditions was used as a positive control [36].

Increased growth in EcN when supplemented with both EA and vitamin B_12_ was observed (Figure 2A-C), indicating catabolism of EA under conditions expected for *eut* BMC utilisation. The absence of increased growth in supplementation with either EA or vitamin B_12_ confirm the vitamin B_12_ dependence of the *eut* BMC in EcN. Low levels of growth are observed even in the absence of a carbon source (Figure 2), as the pre-culture media (M9 containing glucose) was unable to be washed while maintaining the Biosafety Level 2 conditions required for this species.

**Figure 2.**
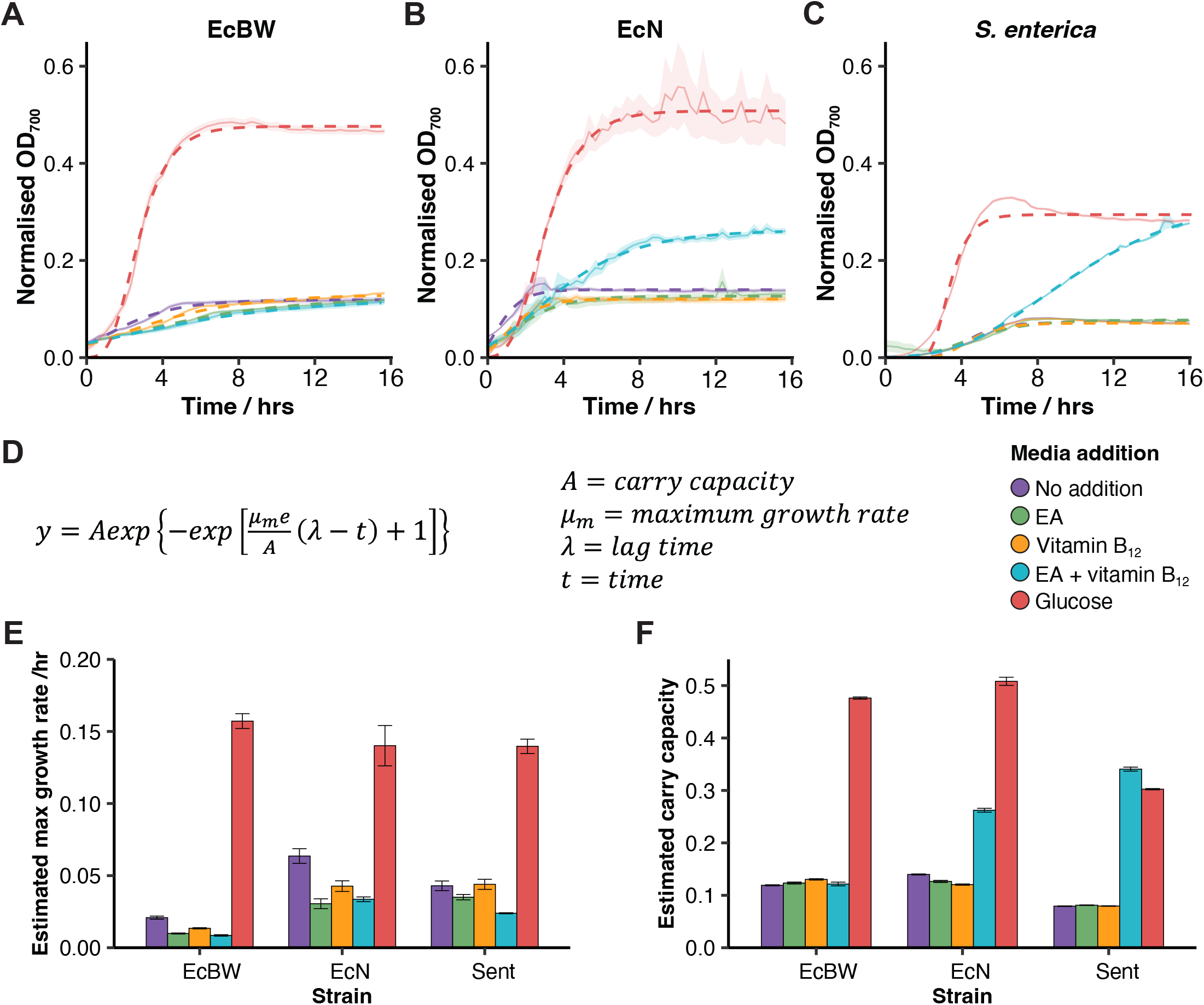
EA-dependent growth in *E. coli* Nissle 1917. Growth curves of EcBW (**A**), EcN (**B**), and *S. enterica* (**C**) when grown in M9 supplemented with either glucose, EA, and/or vitamin B_12_, in triplicate. The mean and standard error are shown with a solid line and a pale ribbon, respectively, and the data was fitted with the Gompertz function (**D**) shown by the respective dashed lines. The estimated growth rates (**E**) and carry capacities (**F**) are shown with standard error bars.

Growth tests supplementing the minimal media with EA, vitamin B_12_ and glucose together were performed to assess the difference in growth cultures compared to glucose alone (SI Figure 1). There was minimal observed difference between the conditions in EcBW, suggesting that the presence of EA and vitamin B_12_ do not have a detrimental or toxic affect to a glucose-metabolising cell. In EcN, the growth with glucose, EA and vitamin B_12_ was higher than when catabolising EA as the sole carbon source (Figure 2B), but lower than glucose alone, suggesting that glucose is both a preferable carbon source and less burdensome than microcompartment formation, as supported by the lower growth rates when utilising EA alone. This also indicates that the expression and utilisation of the *eut* BMC is dependent not only on the presence of both EA and vitamin B_12_, but also the absence of glucose.

The Gompertz model (Figure 2D) was used to estimate the growth rates and carry capacities (maximum cell density) of the cultures. While the estimated growth rates were comparable or lower (Figure 2E), the estimated carry capacity of EcN and *S. enterica* are higher in minimal media supplemented with EA and vitamin B_12_ compared to either alone or no addition (Figure 2F). This suggests that while growth is observed under *eut* BMC utilising conditions, the catabolism of EA is burdensome to the cell. This would support the expression of the *eut* BMC only in the absence of glucose in times of nutrient depletion when EA is an adequate alternative, where EA is not a preferred carbon source, but is an adequate alternative in times of nutrient depletion. While *S. enterica* has the capacity for *de novo* vitamin B_12_ synthesis [49], this is only possible under anaeobic conditions, and so exogenous vitamin B_12_ is supplied under the aerobic conditions in this study.

### *2*.*2 eut* BMC expression in EcN

Next, we explored whether BMC structures could be observed directly using transmission electron microscopy (TEM). BMC structures were observed in EcN when supplemented with EA and vitamin B_12_ (Figure 3A), but not with unsupplemented media, or either EA or vitamin B_12_ alone. This suggests that both EA and vitamin B_12_ are required to induce the production of the BMC structure. Additionally, EcBW did not produce BMCs when supplemented with both EA and vitamin B_12_, indicating that the prophage insertion is sufficient in disrupting the expression of the *eut* operon. Structures appear under the same conditions where growth is demonstrated in media supplemented with EA and vitamin B_12_. The BMC structures are the same approximate size (50-200 nm) and shape as previously reported BMCs in other species and *E. coli* strains, as they had not been previously visualised in EcN [50, 51, 52].

**Figure 3.**
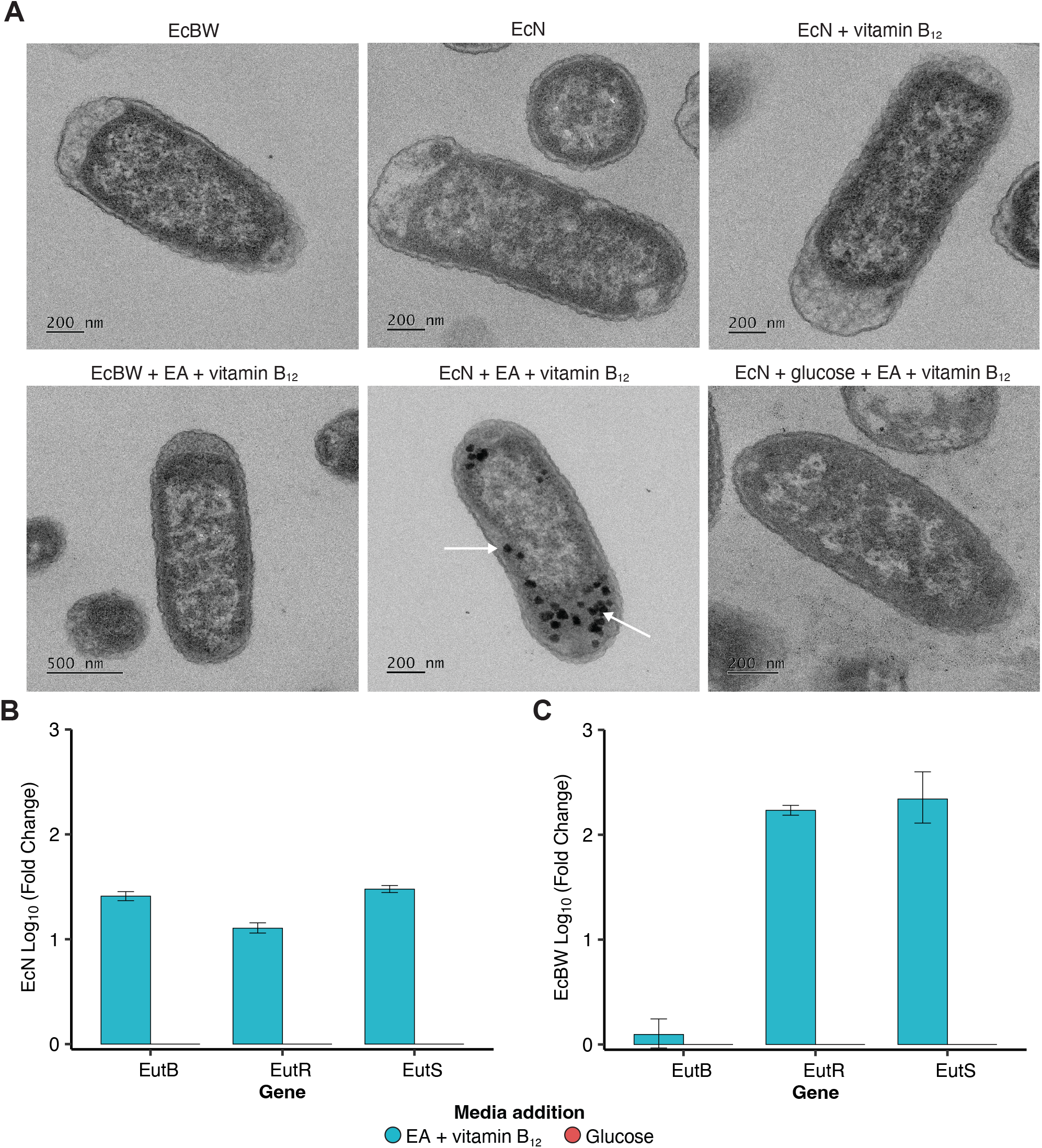
Microcompartment formation in *E. coli* Nissle 1917. (**A**) Transmission electron micrographs of EcBW and EcN transformed with DVA plasmid for antibiotic resistance grown in media supplemented with and without glucose, ethanolamine and vitamin B_12_, where suspected BMC structures are indicated by white arrows. (**B & C**) Change in expression of three key *eut* operon genes in EcN and EcBW, respectively. qPCR was performed in M9 supplemented with either EA and vitamin B_12_, or glucose. The qPCR was completed in both biological and technical triplicate, where the compounded standard error bars are shown.

To further verify the structures observed in the TEM micrographs, qPCR with 3 key *eut* genes was performed under conditions predicted to utilise the *eut* pathway compared to standard glucose metabolism (Figure 3B-C). The gene *eutS* encodes one of the shell proteins, *eutB* encodes part of the signature enzyme for the pathway, and *eutR* is a transcription factor for the operon. As the presence of glucose has previously been reported to supress *eut* expression [43], and this was observed in the growth assays in EcN (Figure 2B), this was a suitable control condition. The mRNA corresponding to the three key *eut* genes had a 12.7 - 30.0 fold increase in expression in EcN when supplemented with EA as the sole carbon source and vitamin B12 as the required cofactor for the *eut* pathway. This result, in combination with the growth assays (Figure 2) and the TEM micrographs (Figure 3A) supports the expression and formation of the *eut* BMC shell, and the expression and utilisation of the enzymes of the *eut* metabolic pathway to utilise EA as a sole carbon source.

Interestingly, the effect of the prophage insertion in the *eut* operon of EcBW in between the genes encoding *eutA* and *eutB* (Figure 1B) is evident from the qPCR results. It was unknown whether the prophage distrupts the expression of the whole operon, or just renders the metabolic pathway non-functional as it interrupts the genes encoding the signature enzyme, preventing EA catabolism. The qPCR results (Figure 3C) indicate that there is increased expression of *eutR* and *eutS* in EA conditions, but not of *eutB*. Despite the shell protein *eutS* being expressed, no BMC structures are observed in identical conditions when imaged with TEM (Figure 3A).

### 2.3 Ammonium production in EcN

Flux balance analysis can be a powerful tool for the exploration and understanding of flux dynamics occurring within a cell given a set of constraints (Figure 4A) [53, 54]. One use for FBA is the generation of experimental hypotheses that can be readily tested *in vitro*. For FBA in EcN we used a modified version of the iHM1533 model [55] (here called iHM1533 CC), which originally predicted that EcN could not catabolise EA as a carbon source, leading to the removal of the EA transport reaction to render the *eut* pathway non-functional by van t’ Hof *et al*. However, since our experimental data demonstrated the ability of EcN to utilise the *eut* pathway (Figures 2 & 3), this transport reaction was restored for this study. Similarly, the iJO1366 model [56] was used for EcBW and predicts EA catabolism, as from the genome-scale model it would appear to express the requisite proteins. However, due to the prophage insertion (Figure 1B), and confirmed by our growth assays (Figure 2A), this is not the case and so the models were modified to reflect this, creating iHM1533 CC and iJO1366 CC (SI tables 3 & 4). The modified models were verified and the flux pathways were visualised (Figure 4B).

**Figure 4.**
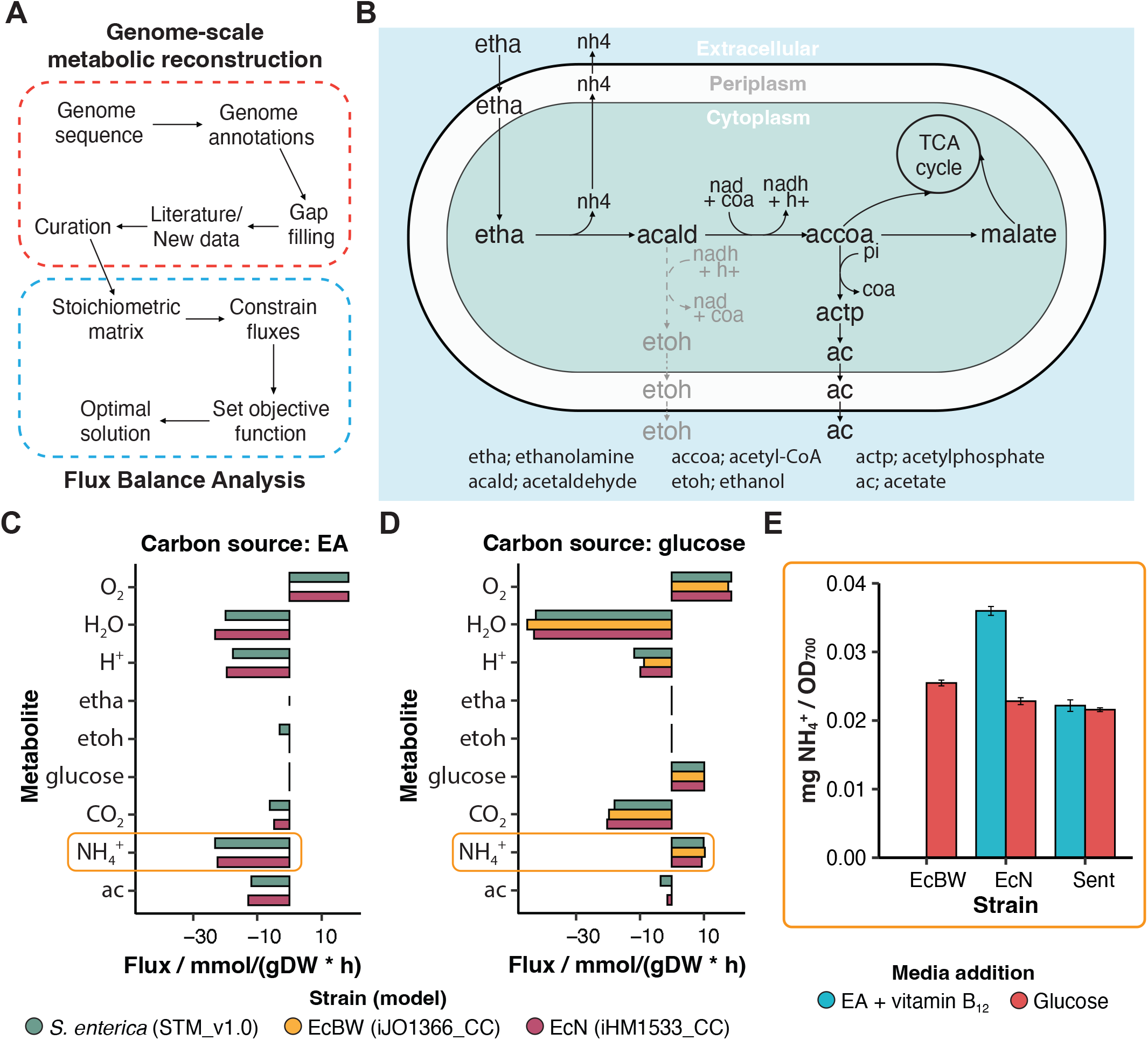
Metabolite flux prediction and experimental verification in *E. coli* Nissle 1917. (**A**) Overview of genome-scale metabolic reconstruction and subsequent flux balance analysis. (**B**) Schematic representing the predicted direction of flux through the *eut* pathway using FBA with iHM1533 CC. The dotted grey line represents pathways without flux. (**C & D**) Predicted flux from the FBA when the model is supplied with either EA or glucose as the sole carbon source, respectively. (**E & F**) 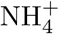 and acetate detected in cell culture after 16/24 hrs growth in media supplemented with either EA and vitamin B_12_, or glucose, in triplicate. The results are normalised to the OD_700_ of the solution, and standard error bars are shown.

Within this study we used FBA models to search for metabolites that would be secreted only if the *eut* pathway was metabolically active, indicating the impact of *eut* utilisation on the environment and wider microbial community. The ammonium generated by the EutBC catabolism of ethanolamine to acetaldehyde can be utilised by the cell as a source of reduced nitrogen for processes such as amino acid synthesis [43, 57]. Many studies report the utilisation of the EA as a primary nitrogen source, where this ammonia generated would produce an additional nitrogen supply to the cell [58, 57]. As expected, the models all performed similarly to each other in FBA in terms of their metabolite uptake and secretion, with the exception of iJO1366 CC which cannot utilise EA (Figure 4C & D, Figure S2). Additionally, the use of EA as a nitrogen source as well as a carbon source can be demonstrated here, where when glucose is supplied as the carbon source, 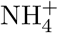 is taken up as a nitrogen source, but when EA is supplied as the carbon source, it is instead secreted (Figure 4C & D). This was expected to be due to the 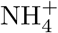 produced when EutBC converts EA into acetaldehyde, and this was confirmed through following the direction of flux in the FBA (Figure 4B). The FBA predicts that the products of *eut* degradation are sequestered directly into the TCA cycle under aerobic conditions, and therefore aid cell growth, and the secretion of 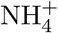 (Figure 4B).

When tested experimentally, it was found that EcN does indeed secrete more ammonium when utilising the *eut* BMC compared to glucose metabolism, but that *S. enterica* does not (Figure 4E). These opposing metabolic outputs have not previously been reported, and while the *eut* pathway is very similar in both species, this highlights a key difference in the surrounding metabolism and adaptations between the two.

### 2.4 Control of EA-dependent growth

In order to explore the potential applications of BMCs within microbial engineering, the expression of the operon needs to be controllable to optimise the formation of the microcompartment and the utilisation of the alternative metabolic pathway it contains. In *E. coli*, 16 of the 17 genes encoded within the *eut* operon are under the inducible control of vitamin B_12_ (Figure 1B). To explore the degree of this control in the *eut*-dependent growth of both EcN and *S. enterica*, growth time courses in minimal media supplemented with EA and varying concentrations of vitamin B_12_ were carried out (Figure 5). In EcN, concentrations of 0 - 32.5 nmol results in a proportional positive effect on growth, with no further increase in growth seen above 40 nmol (Figure 5 A & C). In *S. enterica*, the plataeu is reached at a lower concentration (25-40 nmol) (Figure 5 B & D), but until this point the growth of the bacteria is proportional to the concentration of exogenous vitamin B_12_ supplied. This indicates that the expression of the operon and utilisation of the resulting BMCs are able to be reasonably controlled by exogenous application of vitamin B_12_.

**Figure 5.**
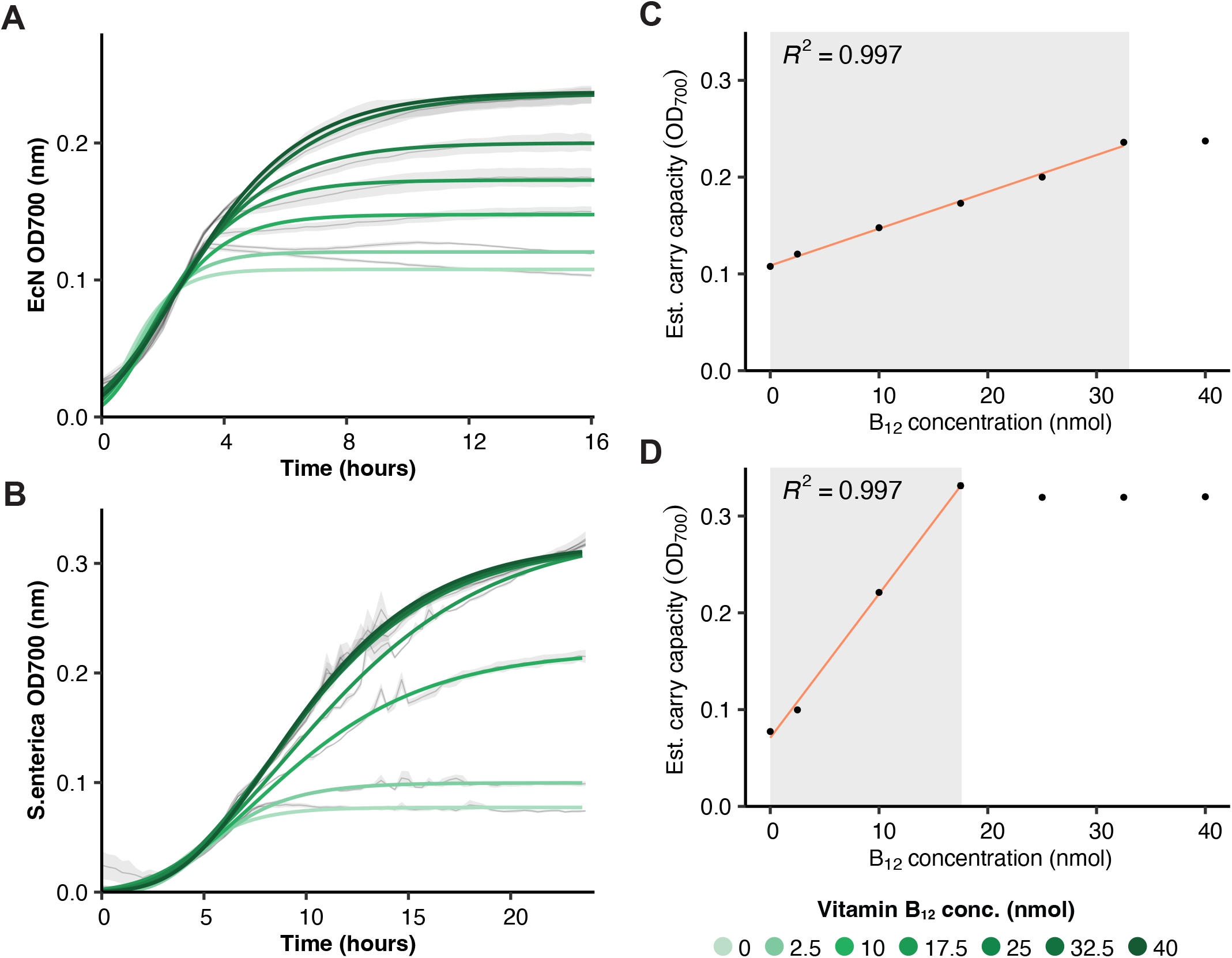
Control of EA-dependent growth in *E. coli* Nissle 1917 and *S. enterica*. (**A & B**) Growth curves of EcN and *S. enterica* when grown in M9 supplemented with EA and different concentrations of vitamin B_12_. The mean and standard error are shown in black and grey, respectively. Gompertz model gits are given by the solid lines. (**C & D**) The estimated carry capacity from the growth curves, with the calculated R^2^ shown above.

## 3 Discussion

Here we demonstrate that EcN shows increased growth when grown in minimal media supplemented with both EA and vitamin B_12_ in the absence of glucose (Figure 2B), when compared to EcBW which cannot utilise the *eut* BMC, and *S. enterica* that can. These results, alongside the TEM micrographs which show BMC-like structures comparable to that of the literature, and the qPCR which showed increased expression of mRNA of three key genes of the *eut* operon, support the hypothesis that EcN can utilise the *eut* BMC to degrade EA not only as a nitrogen source [40], but also as the sole carbon source. Additionally, the expression of the shell protein EutS in EcBW (Figure 3C) but lack of BMC shell formation (Figure 3A) could indicate that the prophage insertion has more widespread effects, for example on the other shell proteins EutL and EutK further downstream. Additionally, the increased secretion of 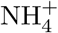 during EA metabolism compared to glucose metabolism in EcN but not in *S. enterica* indicates different adaptations and subsequent metabolism of the products of the *eut* pathway despite the similarities in the *eut* operon and BMC structure between the two. The effects of this difference on supporting growth of the extended microbial community in the gut is not yet known, but an interesting avenue for investigation.

In context of *eut* BMC use in enteropathogens for colonisation advantage, this places EcN within a niche that was previously thought to be predominantly occupied by pathogenic species [18, 32]. Additionally, the secretion of 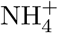 from EA degradation could provide an insight into how EA-degradation interacts with the environment, and perhaps indirectly affects the growth of other populations in the gut microbiome. By understanding how these dynamics work in pathogens and their commensal nutrient-competitors, we can further investigate methods of bacterial community control. This work also shows the control of EA-dependent growth using the native promoter, and lays the groundwork for potential engineering of the promoter to be under the control of an inducer that does not naturally occur in the desired environment. In this way, the growth of EcN could be tunable, and could potentially be controlled to compete with other *eut* BMC utilising bacteria, for example *S. enterica*. Here we demonstrated the ability to control the EA-dependent growth of EcN and *S. enterica* by inducing the *eut* operon.

The FBA was used to compare genome-scale models to predict metabolite uptake and secretion to identify any potential cross-feeding or interaction dynamics that could be exploited for synthetic community design. When the models were updated based on the experimental findings of this study, the FBA was utilised to generate hypotheses regarding metabolite secretion into culture media which were then tested experimentally. Through this method, it was found that EcN secretes increased levels of 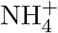 when catabolising EA as opposed to glucose (Figure 4), which could contribute to changes in the ecological landscape of the bacterial community. However, *S. enterica* was not found to increase 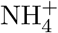 secretion under EA catabolising conditions, perhaps indicating their different effects on the wider gut microbial community.

As FBA does not capture biological dynamics, such as nutrient sensing mechanisms or promoter induction, the results need to be contextualised given literature and experimental findings to interpret them correctly. Additionally, the directive of FBA is to optimise the flux given a set objective function, where this may not accurately capture the true biological mechanisms. For example, the direction of flux through the *eut* pathway predicted by the FBA in this study (Figure 4B) predicts that there will be no flux through the ethanol production branch of the pathway. However, Huseby *et al*. found that the cofactors involved in this reaction, namely the NAD^+^, NADH and CoA, are actually recycled within the compartment, and act to pull the acetaldehyde through the reactions quicker, to convert it into less toxic and volatile forms [24]. This is important because the pores of the microcompartment shell are not selective enough to prevent acetaldehyde escape given sufficient time and concentration. However, while these limitations exist, they are significantly mitigated through critical application and verification in the context of existing literature and new experimental data. In this way, FBA proved a powerful resource in generating hypotheses of metabolite activity to test in the lab, and allowed the rapid assessment and selection of metabolic pathways that were most relevant to our investigation of the effects of *eut* BMC utilisation on the gut microbial community.

This study extends the previous work of this field in characterising the role of BMC-mediated EA utilisation as a carbon source in the absence of glucose in a common gut commensal, *E. coli* strain Nissle 1917. The prevalence of this mechanism in a gut commensal, supporting the work of recent studies [39, 40], highlights that the community dynamics between pathogen and commensal in the gut could be more complex than previously thought, and that there could be competition for a niche that was established as a mechanism for pathogen colonisation. This work contributes to the potential to leverage BMC-dependent growth of a commensal strain to deplete the nutrient pool of a pathogenic strain and thus outcompete it [59, 26]. The difference in ammonium production between EcN and *S. enterica* perhaps suggests different commensal community support to maintain suppression of an pathogen after its been outcompeted, indicating more complex dynamics over this niche than previously thought. As outlined in previous reports and reviews [39, 18], understanding not only the mechanisms of the *eut* BMC, but also this interaction between the gut microbiota and potential pathogens provides many opportunities for gut community control for health, and for microbial engineering applications. This also lays the ground work for the use of the *eut* BMC as a modular metabolic unit within engineering biology, where it could be used to control community dynamics. Additionally, we have shown that FBA can be a useful tool for hypothesis generation when characterising pathways experimentally.

## 4 Methods

### 4.1 Strains

All strains used within this study are listed in Table 1. Ampicillin resistant strains were created from electrotransformation of electrocompetent cells of the relevant strain with the DVA plasmid encoding ampicillin resistance obtained from the CIDAR MoClo kit [60].

**Table 1:** Bacterial strains used in this study.

| Strain | Description | Source |
| --- | --- | --- |
| *Escherichia coli* BW25113 | Assumed BMC negative lab strain | Keio collection [61] |
| *Escherichia coli* Nissle 1917 | Assumed BMC positive commensal | Prof Ian Henderson |
| *Escherichia coli* BW25113 - DVA | Ampicillin resistant lab strain | This study |
| *Escherichia coli* Nissle 1917 - DVA | Ampicillin resistant commensal | This study |
| *Salmonella enterica subsp. enterica* serotype Typhimurium | BMC positive enteropathogen | Leibniz Institute DSMZ |

### 4.2 Culture media and growth conditions

Overnight cultures of BW25113 and EcN were grown at 37^*°*^C with shaking (200 rpm) from single colonies picked from LB agar plates, in 5 mL of supplemented M9 minimal media (0.2% casamino acids, and either 0.4% glycerol (v/v) or 0.4% glucose (v/v)) and 100 μg/mL Ampicillin where appropriate.

The ability of the strains to grow with EA as a sole carbon source was tested on minimal media supplemented with 150 nmol 100% cyanocobalamin as vitamin B_12_ (Merck Life Science Limited, UK) and 30 mmol *≥* 99.0% EA hydrochloride (Sigma-Aldrich, UK), unless specified otherwise.

### 4.3 Growth assays

The overnight cultures were diluted to OD_700_ 0.05 in M9 media containing Ampicillin, EA and vitamin B_12_ as appropriate. Then 125 *μ* L of the diluted culture was added to a 96-well microtitre plate (Greiner, DE) covered with a Breathe-Easy sealing membrane (Diversified Biotech, USA). The plate was inserted into a Tecan Spark plate reader (Tecan, UK), and grown for 16 hours with continuous double orbital shaking (2 mm, 150 rpm) at 37^*°*^C. Absorbance at 700 nm was measured every 20 minutes. Calibration and normalisation of plate reader data was performed in R using the freely available FlopR package [62]. Plots were made in R [63] using ggplot2 [64] and dplyr [65].

Additional growth assays were completed using spectrophotometer (Thermo Scientific Multiskan GO) measurements of the absorbance at 700 nm every 2 hours for 6 hours. The overnight cultures were diluted to OD_700_ 0.05 in M9 media containing Ampicillin, EA and vitamin B_12_ as appropriate. Then 5 mL of the diluted culture was added to 14 mL culture tubes and incubated at 37^*°*^C with shaking (200 rpm) for the duration of the assay.

#### Fitting data

Gompertz model [66, 67] was used in R to fit the cell growth curves (Figures 2 & 5) for each sample with measured cell density (OD_700_). The growth curves were defined as the logarithm of the cell density (y) plotted against time (t):

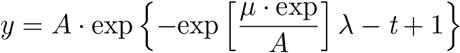

where *μ* stands for the growth rate of the bacteria at exponential growth phase; A is the maximum cell density that could be achieved by the cell culture; *λ* is the lag time before the bacteria enter exponential growth phase.

### 4.4 Transmission Electron Microscopy

#### Culture preparation and cell embedding

After growth (as described above) of 10 mL of each culture was pelleted by centrifugation. The pellet was fixed in 2 mL of 2.5% glutaraldehyde diluted in 100 mM sodium cacodylate, pH 7.2 for 2 hrs with gentle rotation. The bacteria were washed twice with 100 mM sodium cacodylate for 10 mins each. The bacteria were then stained for 2 hrs in 1 mL 1% (w/v) osmium tetroxide diluted in 100 mM sodium cacodylate. The pellets were washed four times with 100 mM sodium cacodylate for 5 mins each. The pellets were dehydrated through an ethanol (EtOH) gradient as follows: 50% (v/v) EtOH for 15 min, 70% EtOH overnight, 90% EtOH for 15 min, and 100% EtOH for 15 min four times. The pellets were then washed twice in propylene oxide for 15 min. The pellets were embedded in 1 mL propylene oxide-low-viscosity (LV) resin (1:1) for 30 min, followed by incubation in fresh LV resin twice for 90 min each. The pellets were resuspended in 1 mL LV resin and transferred too a conical-bottom tube, and then following centrifugation were left to polymerise at 60^*°*^C overnight.

#### Sectioning and visualisation of samples

Samples were ultra-thin sectioned on an RMC MT-XL ultra-microtome with a diamond knife (diatome 45^*°*^) sections (60-70 nm) were collected on un-coated 300 mesh copper grids. Grids were stained by incubation in 4.5% uranyl acetate in 1% acetic acid solution for 45 minutes followed by washing in a stream of dH_2_0. Grids were then stained with Reynolds lead citrate for 7 minutes followed by washing in a stream of dH_2_0.

Electron microscopy was performed using a JEOL-1230 transmission electron microscope equipped with a Gatan multiscan digital camera operated at an accelerating voltage of 80 kV.

### 4.5 Ammonia production assay

The overnight cultures were diluted in appropriate media to OD700 = 0.05, or 1:1000 in accordance to appropriate BioSafety Level conditions. After 16 and 24 hrs growth for *E. coli* strains and *S. enterica*, respectively, to ensure completion of the growth phase, the ammonia detection assay was completed as per manufacturer instructions (Sigma-Aldrich).

### 4.6 qPCR

Total RNA from 16 hour growth cultures (as described above) was extracted using the Qiagen RNeasy Plus kit (Qiagen, Chatsworth, CA). RNA was treated with DNaseI (amplification grade; Invitrogen) to remove any remaining trace amounts of DNA. cDNA was generated with 450 ng of RNA by use of the Qiagen Whole Transcriptome Amplification Kit as described by the manufacturer. Primers for qPCR can be found in Table 2. The qPCR was performed with SYBR Green reagent on a Chromo4 PTC-200 Real-Time PCR Detector system (Bio-Rad). PCR conditions were 94°C for 2 min, followed by 40 three-step cycles of 94°C for 15 s, 60°C for 30 s, and 72°C for 30 s. *rrsA* was used for endogenous reference gene control. Each primer set amplified a single PCR product of predicted size as determined by melt-curve analysis following PCR and by agarose gel electrophoresis, and had approximately equal amplification efficiencies when validated with a serial dilution of representative cDNA. Each qPCR was performed in triplicate, and relative quantification was determined according to the *δδ*C(t) method [68].

**Table 2:**
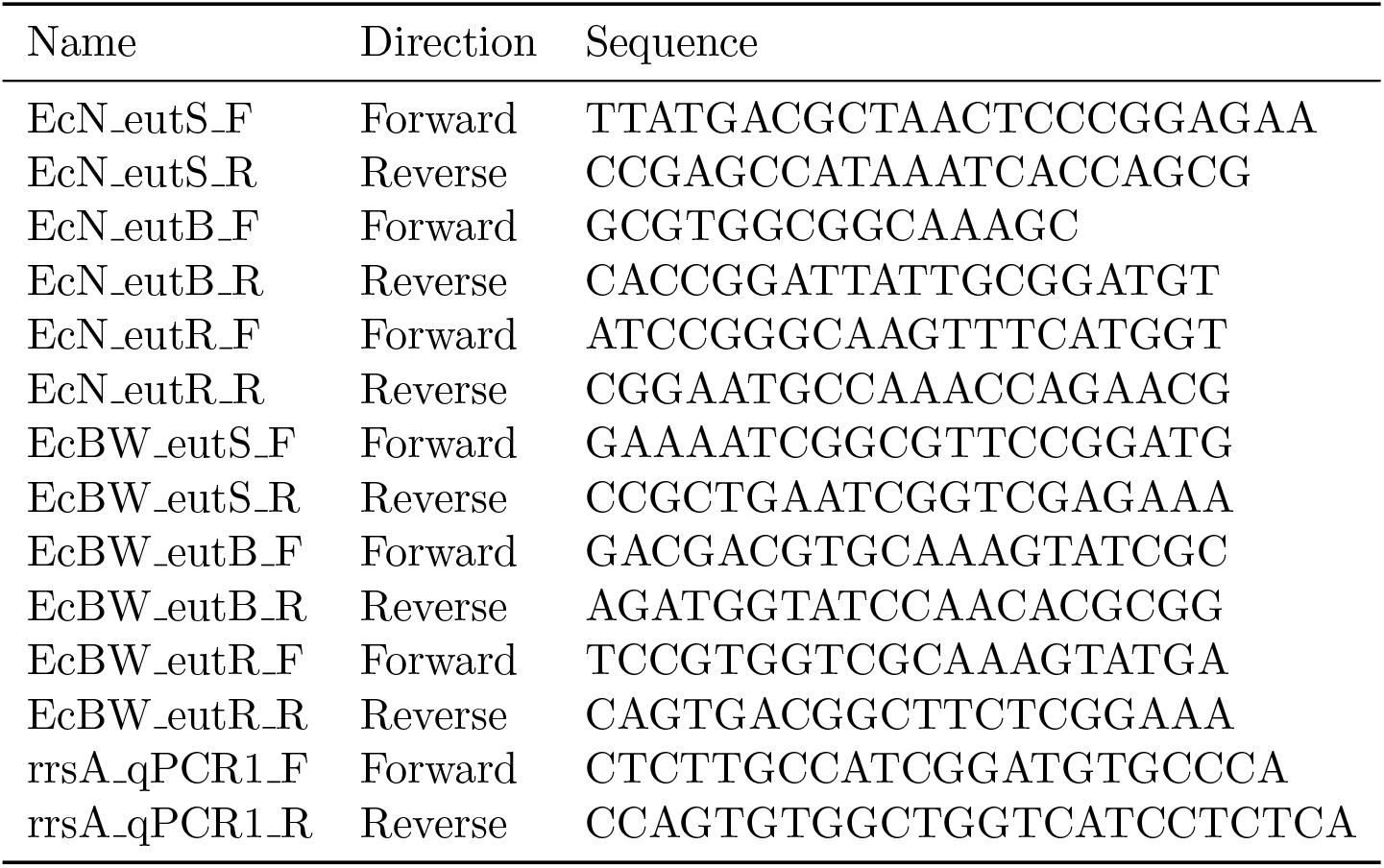
Primers used in qPCR.

### 4.7 Flux Balance Analysis

Flux balance analysis was completed on a MacBook Pro A2338 in both MATLAB using the COBRA Toolbox [69] with Gurobi [70] as the solver, and in Python using the COBRApy Toolbox [71] using Jupyter Notebook. The iHM1433 [55], iJO1366 [56] STM v1.0 [72] models were used for *E. coli* Nissle 1917, *E. coli* BW25113, and *S. enterica*, respectively.

FBA code repository can be found at https://github.com/ChaniaClare/FBA-eut-BMC/

## 5 Acknowledgements

CC was supported by the Biotechnology and Biological Sciences Research Council (grant no. BB/T008709/1). JWR was supported by the Engineering and Physical Sciences Research Council (EPSRC) (grant no. EP/W004674/1). SF was supported by SF was supported by EPSRC (grant no. EP/R013756/1) through the Future Vaccine Manufacturing Research Hub (Vax-Hub). CPB was supported through the European Research Council (ERC) under 254 the European Union’s Horizon 2020 research and innovation programme (grant no. 770835), and by the Wellcome Trust (grant no. 209409/Z/17/Z).

We thank Dr Bill Andrews of the UCL Molecular Biology Facility for completing the qPCR, and Elizabeth Slavik-Smith of the Biosciences EM core facility for her TEM instruction and assistance.

