## Supplementary Information for "Bacterial microcompartment utilisation in the human commensal *Escherichia coli* Nissle 1917"

### 1 Extended materials and methods

Table 1: **Protein sequence homology of proteins comprising *eut* operon between EcN and BW25113**

| Protein | No. amino acids | % identity between EcN and BW25113 |
|---|---|---|
| eutS | 111 | 100.00 |
| eutP | 159 | 97.48 |
| eutQ | 233 | 97.00 |
| eutT | 267 | 100.00 |
| eutD | 338 | 99.70 |
| eutM | 97 | 100.00 |
| eutN | 95 | 100.00 |
| eutE | 467 | 98.93 |
| eutJ | 278 | 98.56 |
| eutG | 395 | 98.73 |
| eutH | 408 | 99.02 |
| eutA | 467 | 98.07 |
| eutB | 453 | 99.78 |
| eutC | 295 | 99.66 |
| eutL | 219 | 97.26 |
| eutK | 166 | 98.80 |
| eutR | 350 | 98.86 |

Table 2: **Summary of each model used in this study**

| | **iJO1366** BW25113 2011 | **iHM1533** EcN 2022 | **STM_v1.0** Sent 2011 |
|---|---|---|---|
| Chromosome size of strain (Mb) | 4.63 | 5.44 | 4.86 |
| Number of reactions | 2583 | 3143 | 2546 |
| Number of metabolites | 1805 | 2261 | 1802 |
| Number of genes | 1367 | 1533 | 1271 |
| Compartments | cytosol; extracellular space; periplasm | cytosol; extracellular space; periplasm | cytosol; extracellular space; periplasm |

Table 3: **Inclusion of metabolites of the *eut* BMC metabolic pathway within each model.** Metabolite ID and names are given as standard GEM notation, and each model name is given alongside its species and year the model was created.

| Metabolite Name | Metabolite ID | iJO1366 EcBW 2011 | iJO1366_CC EcBW 2024 | iHM1533 EcN 2022 | iHM1533_CC EcN 2024 | STM v1.0 Sent 2011 |
|---|---|---|---|---|---|---|
| Acetaldehyde | acald_e | × | × | × | × | × |
| Acetaldehyde | acald_c | ✓ | ✓ | ✓ | ✓ | ✓ |
| Acetyl phosphate | actp_c | ✓ | ✓ | ✓ | ✓ | ✓ |
| Acetyl-CoA | accoa_c | ✓ | ✓ | ✓ | ✓ | ✓ |
| Ammonium | nh4_e | ✓ | ✓ | ✓ | ✓ | ✓ |
| Ammonium | nh4_p | ✓ | ✓ | ✓ | ✓ | ✓ |
| Ammonium | nh4_c | ✓ | ✓ | ✓ | ✓ | ✓ |
| Coenzyme A | coa_c | ✓ | ✓ | ✓ | ✓ | ✓ |
| Ethanol | etoh_e | ✓ | ✓ | ✓ | ✓ | ✓ |
| Ethanol | etoh_p | ✓ | ✓ | ✓ | ✓ | ✓ |
| Ethanol | etoh_c | ✓ | ✓ | ✓ | ✓ | ✓ |
| Ethanolamine | etha_e | ✓ | ✓ | ✓ | ✓ | ✓ |
| Ethanolamine | etha_p | ✓ | ✓ | ✓ | ✓ | ✓ |
| Ethanolamine | etha_c | ✓ | ✓ | ✓ | ✓ | ✓ |
| H+ | h_e | ✓ | ✓ | ✓ | ✓ | ✓ |
| H+ | h_p | ✓ | ✓ | ✓ | ✓ | ✓ |
| H+ | h_c | ✓ | ✓ | ✓ | ✓ | ✓ |
| NAD | nad_c | ✓ | ✓ | ✓ | ✓ | ✓ |
| NADH | nadh_c | ✓ | ✓ | ✓ | ✓ | ✓ |
| Phosphate | pi_e | ✓ | ✓ | ✓ | ✓ | ✓ |
| Phosphate | pi_p | ✓ | ✓ | ✓ | ✓ | ✓ |
| Phosphate | pi_c | ✓ | ✓ | ✓ | ✓ | ✓ |

Table 4: **Inclusion of reactions of the *eut* BMC metabolic pathway within each model.** Reaction ID and names are given as standard GEM notation, and each model name is given alongside its species and year the model was created.

| Reaction ID | Reaction | iJO1366 EcBW 2011 | iJO1366_CC EcBW 2024 | iHM1533 EcN 2022 | iHM1533_CC EcN 2024 | STM v1.0 Sent 2011 |
|---|---|---|---|---|---|---|
| Ex_acald_e | acald_e <=> | ✓ | ✓ | ✓ | ✓ | ✓ |
| ALCD2x | etoh_c + nad_c <=>acald_c + h_c + nadh_c | ✓ | ✓ | ✓ | ✓ | ✓ |
| EX_nh4_e | nh4_e <=> | ✓ | ✓ | ✓ | ✓ | ✓ |
| NH4tpp | nh4_p <=>nh4_c | ✓ | ✓ | ✓ | ✓ | ✓ |
| NH4tex | nh4_e <=>nh4_p | ✓ | ✓ | ✓ | ✓ | ✓ |
| Ex_etoh_e | etoh_e <=> | ✓ | ✓ | ✓ | ✓ | ✓ |
| ETOHtrpp | etoh_p <=>etoh_c | ✓ | ✓ | ✓ | ✓ | ✓ |
| ETOHtex | etoh_e <=>etoh_p | ✓ | ✓ | ✓ | ✓ | ✓ |
| ETHAAL | etha_c –>acald_c + nh4_c | ✓ | ✓ | ✓ | ✓ | ✓ |
| EX_etha_e | etha_e –> | ✓ | ✓ | ✓ | ✓ | ✓ |
| ETHAt2pp | etha_p + h_p –>etha_c + h_c | ✓ | ✓ | ✓ | ✓ | ✓ |
| ETHAtex | etha_e <=>etha_p | ✓ | × | × | ✓ | ✓ |
| Ex_h_e | h_e <=> | ✓ | ✓ | ✓ | ✓ | ✓ |
| Ex_pi_e | pi_e <=> | ✓ | ✓ | ✓ | ✓ | ✓ |
| PItex | pi_e <=>pi_p | ✓ | ✓ | ✓ | ✓ | ✓ |
| PTAr | accoa_c + pi_c <=>actp_c + coa_c | ✓ | ✓ | ✓ | ✓ | ✓ |
| Htex | h_e <=>h_p | ✓ | ✓ | ✓ | ✓ | ✓ |

### 2 Extended results

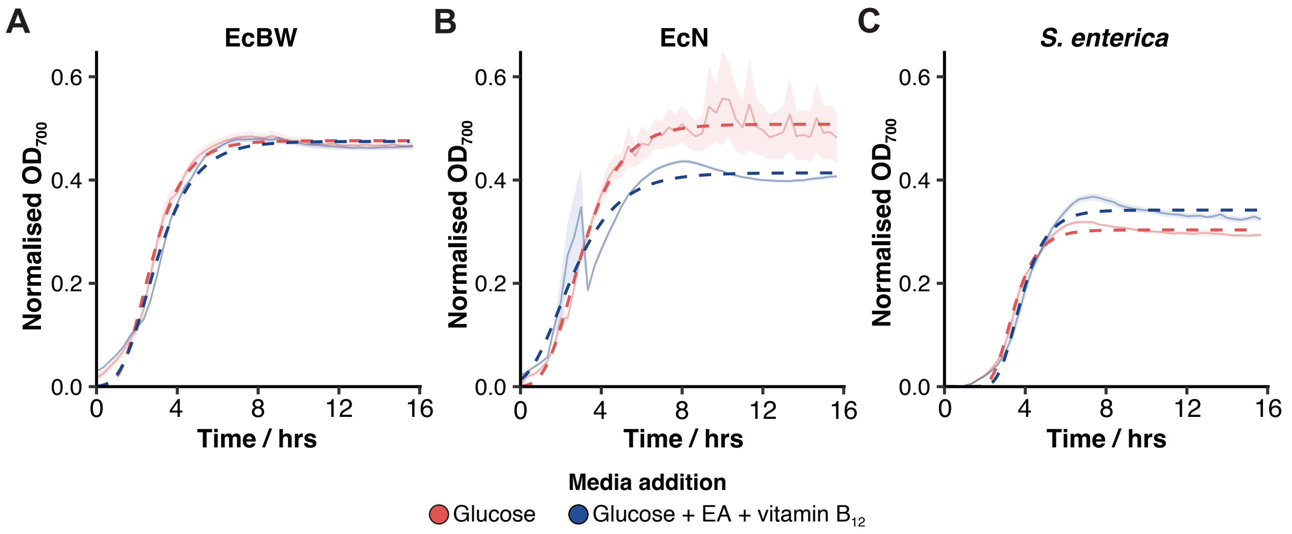


Figure 1: **EA-dependent growth in *E. coli* Nissle 1917.** Growth curves of EcBW (**A**), EcN (**B**), and *S. enterica* (**C**) when grown in M9 supplemented with either glucose alone, or EA, vitamin $B_{12}$ and glucose, in triplicate. The mean and standard error are shown with a solid line and a pale ribbon, respectively, and the data was fitted with the Gompertz function shown by the respective dashed lines.

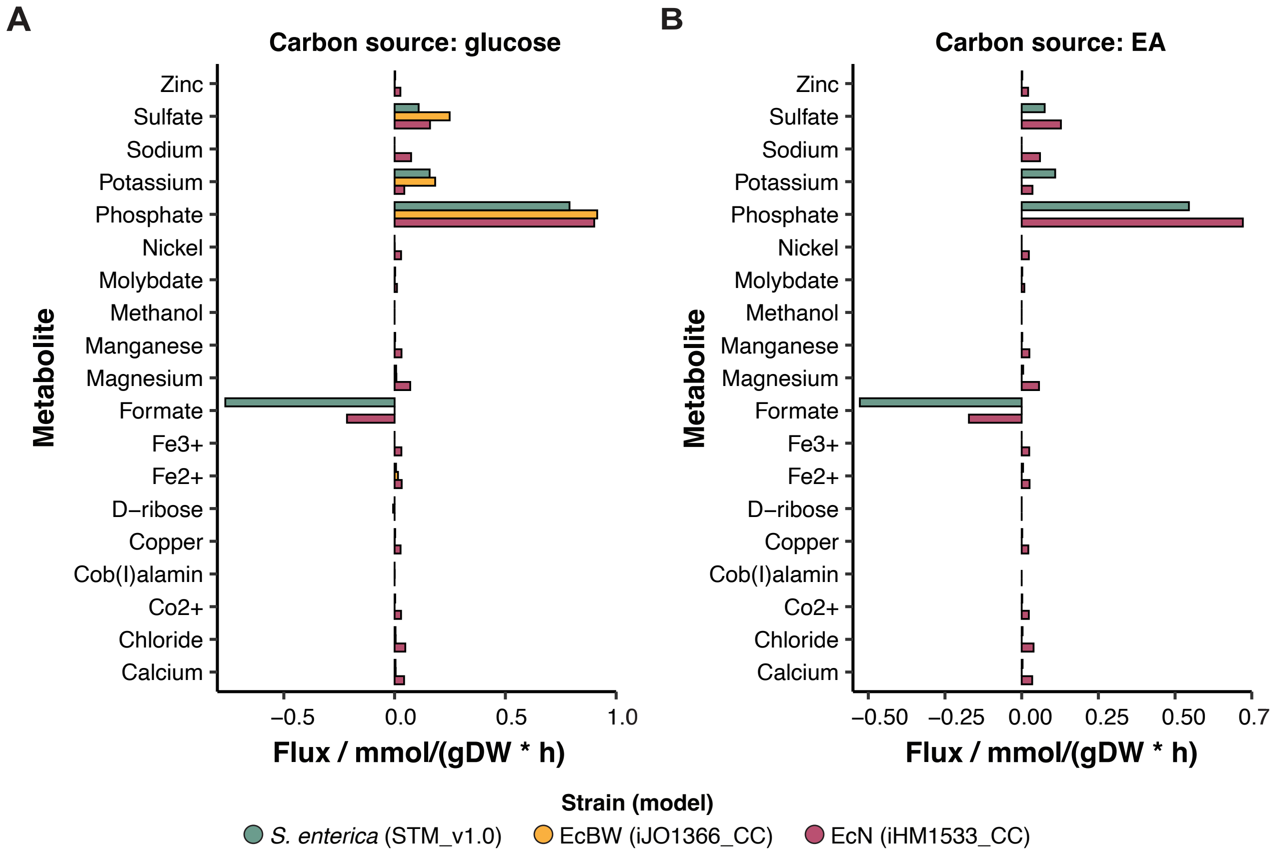


Figure 2: **Metabolite flux prediction in *E. coli* Nissle 1917.** (**A & B**) Predicted flux from the FBA when the model is supplied with either EA or glucose as the sole carbon source, respectively .
